# Biological and structural analysis of new potent Integrase-LEDGF allosteric HIV-1 inhibitors

**DOI:** 10.1101/2023.01.28.523533

**Authors:** Erwann Le Rouzic, Damien Bonnard, Frédéric Le Strat, Claire Batisse, Julien Batisse, Céline Amadori, Sophie Chasset, Stéphane Emiliani, Benoit Ledoussal, Marc Ruff, François Moreau, Richard Benarous

## Abstract

LEDGF/p75 (LEDGF) main cellular cofactor of HIV-1 integrase (IN), acts as tethering factor to target integration of HIV in actively transcribed genes. Recently a class of IN inhibitors based on inhibition of LEDGF-IN interaction has been developed. We describe here a new series of IN-LEDGF allosteric inhibitors (INLAIs), with potent anti-HIV-1 activity in the one-digit nanomolar range. These compounds inhibited IN-LEDGF interaction while enhancing IN-IN aberrant multimerization by allosteric mechanism. Compounds of this series were fully active on HIV-1 mutants resistant to IN strand transfer inhibitors (INSTIs) and other class of anti-HIV drugs, confirming that they belong to a new class of antiretrovirals. These compounds displayed most potent antiretroviral activity at post-integration due to aberrant IN polymerization responsible of infectivity defect of viral particles produced. INLAI-resistant mutants were selected by dose escalation method and the most detrimental mutation found was T174I. The impact of these resistance mutations was analyzed by fold shift EC_50_ with regard to wild type virus and the replication capacity of mutated viruses were determined. Crystal structure of BDM-2, the lead compound of this series in complex with IN catalytic core domain (CCD) was elucidated. No antagonism was observed between BDM-2 and a panel of 16 antiretroviral drugs from different classes. In conclusion, the overall virologic profile of BDM-2 warrants the recently completed single ascending dose phase I trial (ClinicalTrials.gov Id: NCT03634085) and supports further clinical investigation for potential use in combination with other antiretroviral drugs.

**Author summary:** Integrase-LEDGF allosteric inhibitors are a new class of antiretrovirals recently developed that target the interaction of HIV-1 Integrase with its cellular cofactor LEDGF/p75 required for HIV-1 integration in actively transcribed genes. The great interest of developing such new class of antiretroviral is to add to the anti-HIV drug arsenal compounds that are fully active on resistant viruses to all other classes of drugs currently used in clinic. However, none of these inhibitors are currently in clinical use or in late clinical trials. Here we describe a series of highly potent Integrase allosteric inhibitors that can be considered as precursors of a new class of antiretroviral drugs, with fully conserved activity on viruses resistant to currently used anti-HIV drugs, and a lead compound that has no antagonism with a large panel of present anti-HIV drugs and that was successfully validated in a phase I clinical trial.

## Introduction

In 2007 Raltegravir (RAL, Merck), and in 2012 Elvitegravir (EVG, Gilead) first generation of Integrase Strand Transfer Inhibitors (INSTIs) were introduced in the therapeutic arsenal against HIV-1 [1]. In 2013, a second generation of INSTIs with Dolutegravir (DTG, GSK) and more recently Bictegravir (Gilead), have been introduced with great success for the treatment of HIV-1 infection [2,3]. These second generation INSTIs have the great advantage of an improved profile of resistance compared to the first generation of INSTIs. However, viruses resistant to these second generation compounds have been found, in particular in patients with a long history of multiple anti-HIV-1 treatment that suffer multi-resistance to antiretrovirals (ARV) [4,5]. It is thus of paramount importance to offer to these patients alternative treatments with a new class of ARVs fully active against resistant viruses to all drugs currently used in clinic. Very active research is ongoing for the development of new classes of ARV with novel mechanism of action, that are currently in clinical investigation such as maturation inhibitors [6], or recently approved by FDA such as attachment inhibitors [7]. Among these new classes of ARV, Integrase LEDGF Allosteric Inhibitors (INLAIs) [8] alternatively named LEDGINs [9], ALLINIs [10] NCINIs [11–13] or MINIs [10,14] have been developed ([15], and for review see [16]). By disruption of the interaction between HIV-1 Integrase (IN) and its major cellular co-factor LEDGF/p75 [17–21], these compounds inhibit integrase by an alternative allosteric mechanism. By binding to the LEDGF/p75-binding pocket, INLAIs promote a dual anti-HIV-1 activity, with inhibition of HIV-1 integration at actively transcription sites that depends on LEDGF/p75 tethering of IN, and at post-integration step with production of non-infectious progeny virions defective for reverse transcription [17,22–25]. This defect in reverse transcription is due to aberrant IN multimerization during viral particle assembly, that results from INLAIs binding to the LEDGF binding pocket on IN [23–25]. IN binds the viral RNA genome and is essential during virion morphogenesis [26,27]. The INLAI-triggered aberrant IN multimerization does not influence packaging of a functional RNA genome [28] but precludes the incorporation of the viral RNA complex into the capsid core of the assembling particle [25]. This post-integration ARV activity is more potent than that at integration, which is the case for all compounds of this class [8,13,17].

Since the first NCINI and LEDGINs reported by Boehringer Ingelheim (BI) [11] and Zeger Debyser’s laboratory [9], several compounds of this class have been described by pharmaceutical companies and academic laboratories including us [8–16].

In this paper we describe a new series of highly potent INLAI compounds that inhibit lab-adapted HXB2 or NL4-3 HIV-1 replication at single-digit nanomolar concentration and that expand the chemical diversity of this class of HIV-1 inhibitors. A compound of this class has been co-crystallized with IN-CCD and more importantly all these new compounds have been co-crystallized with a construct between integrase CTD and CCD domains. INLAI resistant mutants have also been characterized and no antagonism of these compounds with several anti-HIV drugs used in clinic has been found.

## Results

### Chemical structures of INLAI compounds of this series

The structures of some previously described INLAIs are shown in Fig 1A. Different scaffolds, quinoline for BI-224436 [12], thiophene for MUT-A [29], benzene for Shionogi S-I-82 [30], benzothiazole for GS-9822 [31], pyrrolopyridine for STP0404 [15], have been used for the development of these compounds. However, all these compounds share the same active group characterized by a carboxylic acid linked to a *tert*-butoxy moiety. Biodim has developed novel potent INLAI compounds, MUT713 also called BDM-2, the lead compound of this series, MUT871, MUT872, MUT884 and MUT916 that are investigated in this paper (Fig 1B). These compounds are based on a benzene scaffold and contain either the carboxyl and *tert-butoxy* groups (BDM-2, MUT871) essential for potency in previously described INLAIs, or for MUT872, MUT884 and MUT916 compounds, a derivative of this motif in which the *tert*-butoxy is replaced by a cyclopropyloxy, the carboxyl function being conserved. Compound MUT871 differs from BDM-2 by a methyl substituent on the chromane group.

**Fig 1.**
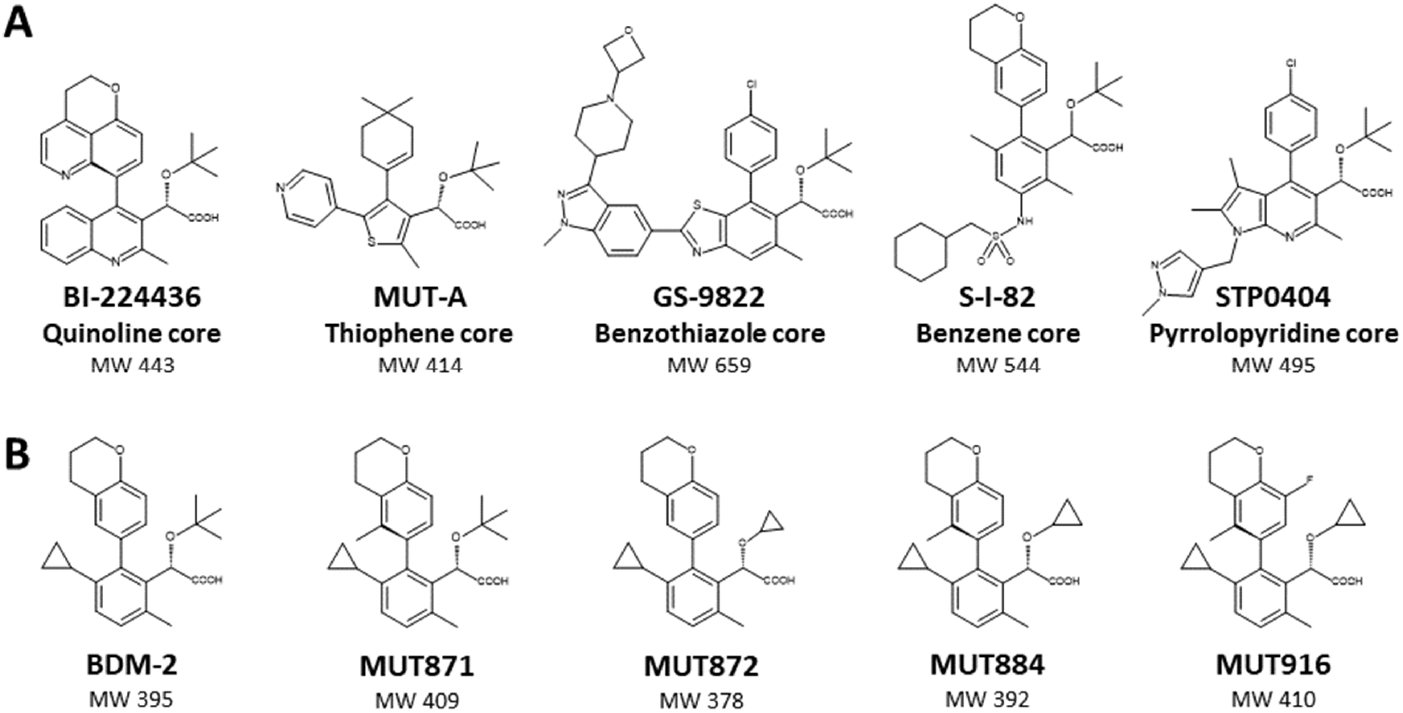
Chemical structure diversity of INLAIs. (A) Previously reported INLAIs are built upon different scaffolds as indicated. All these compounds share the same active group characterized by a carboxylic acid linked to a *tert-butoxy* moiety. (B) Lead compound BDM-2 series. These molecules have a benzene scaffold and contain either a *tert*-butoxy (BDM-2, MUT871), or a cyclopropyloxy group (MUT872, MUT884 and MUT916). MW, molecular weight.

### INLAIs of the BDM-2 series inhibit IN-LEDGF/P75 interaction and promote aberrant IN multimerization

INLAI compounds from this series were tested for inhibition of IN-LEDGF/p75 interaction either between full length proteins or between the binding domains of both proteins, the catalytic core domain of IN (IN-CCD) and the Integrase binding domain of LEDGF/p75 (IBD), using homogeneous time-resolved fluorescence (HTRF^®^) assays (Table 1). The most potent inhibitor of IN-LEDGF/p75 interaction of all INLAIs tested in these assays was MUT871 with 14 nM IC_50_, compared to 90 nM and 82 nM in this assay for previously described inhibitors of this class BI-224436 and S-I-82 respectively. All the other compounds of the BDM-2 series, MUT872, MUT884, MUT916 and BDM-2, are also very potent in disrupting the complex IN-LEDGF/p75 with IC_50_ in the range between 17 nM for MUT872 and 62 nM for MUT884. These compounds were also very active in promoting aberrant IN multimerization, with 50% of maximum multimerization activation (AC_50_) obtained in the same concentration range as for the inhibition of IN-LEDGF/p75, with AC_50_ of 20 nM for BDM-2, and a maximum of 43 nM for MUT872, compared to 34 nM for BI-224436 and 47 nM for S-I-82 in this assay. The plateau reached in this assay means the maximum IN multimerization obtained upon saturation binding of INLAI compounds in the LEDGF binding pocket, compared to the basal IN multimerization measured in the absence of compound. This plateau is between 3.7 to 4 times the basal level for MUT916 and MUT884 respectively to 5-6 times a maximum for BDM-2, in the same range as BI-224436 and S-I-82.

**Table 1.** Biochemical activities of BDM-2 series compounds in IN-CCD– LEDGF/p75-IBD, IN–LEDGF/p75 and multimerization HTRF assays.

| Compound | IN-CCD–<br>LEDGF/p75- IBD<br>IC<sub>50</sub> (μM) | IN–LEDGF/p75<br>IC<sub>50</sub> (μM) | IN multimerization<br>AC<sub>50</sub> (μM)<br>Plateau (%) |
| --- | --- | --- | --- |
| BI-224436 | 0.073 ± 0.013 | 0.090 ± 0.019 | 0.034 ± 0.011<br>670 ± 210% |
| S-I-82 | 0.43 ± 0.13 | 0.82 ± 0.38 | 0.047 ± 0.031<br>500 ± 43% |
| BDM-2 | 0.15 ± 0.03 | 0.047 ± 0.014 | 0.020 ± 0.011<br>560 ± 150% |
| MUT148871 | 0.032 ± 0.014 | 0.014 ± 0.005 | 0.031 ± 0.020<br>440 ± 53% |
| MUT148872 | 0.42 ± 0.07 | 0.17 ± 0.02 | 0.043 ± 0.011<br>440 ± 48% |
| MUT148884 | 0.082 ± 0.021 | 0.062 ± 0.009 | 0.022 ± 0.004<br>400 ± 71% |
| MUT148916 | 0.019 ± 0.008 | 0.046 ± 0.016 | 0.022 ± 0.002<br>370 ± 60% |

Determination of compound IC<sub>50</sub>s for interaction inhibition and for activity to promote IN multimerization with maximum multimerization reached (Plateau) and with compound concentration required for 50% maximum activation of multimerization (AC<sub>50</sub>). Data are mean ± SD of >3 independent experiments.

### INLAI compounds from BDM-2 series exhibit potent antiretroviral activity

Anti-Retroviral (ARV) activity of BDM-2 series was first evaluated in lymphoblastoid cell line MT4 and in primary human T lymphocytes using both NL4-3 and HXB2 laboratory HIV-1 isolates and was compared to that of some INSTIs (Raltegravir (RAL) and Dolutegravir (DTG)) and previously described INLAIs, BI-224436 and S-I-82 (Table 2). ARV activities of BDM-2 and MUT871 were the most potent INLAIs of this series, in the single digit nanomolar range EC_50_ with 8.7 nM and 4.5 nM against NL4-3 and 3.1 nM and 1.4 nM against HXB2 respectively. The other compounds of the BDM-2 series were almost as potent, in the EC_50_ range of 6.3 nM and 8.8 nM for MUT884 and MUT916 respectively on HXB2 while these two compounds were less potent on HIV-1 NL4-3 (EC_50_ of 15 nM and 20 nM respectively). MUT872 was the least potent compound with EC_50_ of 18 nM and 45 nM for infection with HXB2 and NL4-3 respectively. In the absence of human serum, BDM-2 and MUT871 were more potent than RAL and MUT871 was in the same magnitude than that of DTG (EC_50_ of 1.9 nM and 2.7 nM against HXB2 and NL4-3 respectively). All BDM-2 compounds had also more potent ARV than the previously reported INLAI BI-224436, while S-I-82 had an ARV potency similar to that of BDM-2. All BDM-2 compounds had a favorable cytotoxicity profile with CC_50_ between 46 μM and 139 μM, corresponding to very high selectivity index.

**Table 2.** Antiretroviral activity of BDM-2 series on MT4 cells infected by HXB2 or NL4-3 lab-adapted HIV-1 and cytotoxicity selectivity index.

| Compound | ARV activity in MR infection assays in MT4 cells |  |  |  |  | Cytotoxicity | Selectivity index |
| --- | --- | --- | --- | --- | --- | --- | --- |
|  | EC<sub>50</sub><br>(nM)<br>HXB2 | EC<sub>50</sub><br>(nM)<br>NL4-3 | EC<sub>90</sub><br>(nM)<br>NL4-3 | PA-EC<sub>90</sub><br>(nM)<br>NL4-3 | PA-EC<sub>90</sub><br>(ng/mL)<br>NL4-3 | CC<sub>50</sub><br>(μM) | CC<sub>50</sub>/EC<sub>50</sub><br>NL4-3 |
| **RAL** | 8.7 ± 6.0 | 14 ± 9 | 28 ± 14 | 105 ± 38 | 46 ± 17 | 438 ± 29 | 31285 |
| **DTG** | 1.9 ± 1.0 | 2.7 ± 1.3 | 6.3 ± 4.4 | 159 ± 144 | 67 ± 60 | 51 ± 28 | 18888 |
| **BI-224436** | 23 ± 8.4 | 51 ± 17 | 77 ± 34 | 273 ± 103 | 121 ± 46 | >200 | >3921 |
| **S-I-82** | 4.1 ± 0.7 | 12 ± 3 | 21 ± 7.2 | NT | NT | 28 ± 3 | 2333 |
| **BDM-2** | 4.5 ± 1.2 | 8.7 ± 2.8 | 13 ± 5.1 | 127 ± 53 | 50 ± 21 | 69 ± 8 | 7931 |
| **MUT871** | 1.4 ± 0.3 | 3.1 ± 1.0 | 4.6 ± 1.6 | 98 ± 31 | 40 ± 13 | 46 ± 11 | 14838 |
| **MUT872** | 18 ± 1 | 45 ± 12 | 61 ± 19 | 999 ± 373 | 378 ± 141 | 139 ± 29 | 3088 |
| **MUT884** | 6.3 ± 1.1 | 15 ± 5 | 23 ± 13 | 439 ± 124 | 172 ± 49 | 78 ± 12 | 5200 |
| **MUT916** | 8.8 ± 0.8 | 20 ± 9 | 30 ± 15 | 865 ± 418 | 355 ± 172 | 78 ± 6 | 3900 |

Evaluation of ARV activity by EC<sub>50</sub> and Protein-adjusted EC<sub>90</sub> (PA-EC<sub>90</sub>), determination of cytotoxicity (CC<sub>50</sub>) on MT4 cells, and selective index (CC<sub>50</sub>/EC<sub>50</sub> ratio); comparison between two INSTIs RAL and DTG, and two other INLAs, BI-224436 and S-I-82 used as references. Data are mean ± SD of >5 independent experiments.

From the different dose response curves, we inferred the EC_90_ value for each compound (see on Table 2). The human serum effect on the ARV potency of these compounds was determined by extrapolation of the EC_90_ at 100% human serum. This value corresponding to the protein-adjusted EC_90_ (PA-EC_90_), was estimated from experimental data on the compound ARV activities in MT4/NL4-3 infection assays in the presence of 10-50% human serum. As indicated on Table 2, the PA-EC_90_ of BDM-2 was evaluated at 127 nM, corresponding to 50 ng/mL, while MUT871 had a 98 nM PA-EC_90_ i.e., 40 ng/mL. PA-EC_90_ is an important parameter that is used to estimate the clinical inhibitory quotient (IQ) to guide the selection of human dose of drug candidates. These BDM-2 PA-EC_90_ values are of the same order of that of DTG [2] and are more favorable than that of BI-224436 previously reported INLAI.

BDM-2, MUT871 and MUT884 compounds exhibit also potent ARV effect on primary CD4+ T lymphocytes (prepared from peripheral blood lymphocytes (PBL) of normal donors) infected with NL4-3 or HXB-2 lab adapted HIV-1 or with various HIV-1 clinical isolates. As indicated on Table 3, these compounds conserved potent ARV activities on these HIV-1 infected primary lymphocytes, including when clinical isolates were used for infection. ARV potency of BDM-2 was higher than that of BI-224436 or RAL and similar to that of S-I-82. ARV potency of MUT871 was the highest of all INLAIs, not far from that of DTG.

**Table 3.** Antiretroviral activity of BDM-2, MUT871 and MUT884 as representatives of the BDM-2 compound series, in primary T CD4+ lymphocytes infected by lab-adapted HXB2, NL4-3, and HIV-1 clinical isolates.

| Compound | Multiple-round infection assays in primary T CD4+ lymphocytes: EC<sub>50</sub> (nM) |  |  |  |  |  |
| --- | --- | --- | --- | --- | --- | --- |
|  | NL4-3 (B) | HXB2 (B) | #KER2008 (A) | #33913N (B) | #NP1538 (B) | #NP1525 (CRF01_AE) |
| **RAL** | 4.9 ± 2.3 | 2.6 ± 1.4 | 6.7 ± 4.0 | 2.1 ± 0.1 | 1.7 ± 1.9 | 4.0 ± 1.7 |
| **DTG** | 0.53 ± 0.16 | 0.17 ± 0.01 | 0.40 ± 0.21 | 0.35 ± 0.42 | 0.29 ± 0.13 | 0.21 ± 0.09 |
| **BI-224436** | 100 ± 80 | 14 ± 7 | 37 ± 12 | 20 ± 9 | 66 ± 42 | 61 ± 35 |
| **S-I-82** | 9.9 ± 1.1 | 5.2 ± 3.9 | 11 ± 2 | 4.4 ± 1.9 | 8.6 ± 0.5 | 10 ± 4 |
| **BDM-2** | 8.3 ± 4.3 | 3.4 ± 1.0 | 8.1 ± 1.3 | 4.2 ± 1.6 | 13 ± 6 | 11 ± 5 |
| **MUT871** | 2.8 ± 0.5 | 1.8 ± 0.3 | 6.4 ± 1.3 | 1.5 ± 0.6 | 4.0 ± 2.3 | 3.5 ± 0.6 |
| **MUT884** | 15 ± 4 | 10 ± 6 | 19 ± 3 | 5.8 ± 3.2 | 41 ± 2 | 12 ± 3 |

ARV activity was determined by EC<sub>50</sub>. The clade of each virus is indicated in parenthesis; comparison with two INSTIs, RAL and DTG, and with two other INLAIs, BI-224436 and S-I-82. Data are mean ± SD of >5 independent experiments.

### Dual antiretroviral activity of INLAI compounds at integration and post-integration steps evaluated by single round and multiple round HIV-1 infection assays respectively

We investigated the global ARV activity of the compounds of the BDM-2 series using both single round (SR) and multiple round (MR) infection assays. SR infection assays reflect the ARV activity of a compound during an early step of the HIV replication cycle (up to integration) using replication defective HIV-1 pseudotyped virus, and MR infection assays reflect global ARV activity during full replication cycle including post-integration steps, using a fully replicative HIV-1 virus. As shown on Table 4, INSTIs like RAL and DTG used as references here, are as potent in their ARV activity in SR assays than in MR assays, with similar EC_50_ values in both assays. In sharp contrast, BDM-2, MUT871 and of all the other INLAIs tested in MR assays are much more potent than in SR assays. EC_50_s measured in SR assays are in the micromolar range, between 0.63 μM and 9.2 μM for all INLAIs MUT. EC_50_s of INLAIs measured in MR assays are in the nanomolar range with 3.1 nM for the best compound MUT871 and 51 nM for the least active, BI-224436 (Table 4). These results confirmed previous observations, indicating that post-integration inhibition of the HIV-1 replication cycle is the major mechanism contributing to global ARV activity of INLAIs. Interestingly, results on Table 4 show that the EC_50_ ratio between SR and MR varied considerably between different INLAI compounds, from 27 for BI-224436 to 767 for S-I-82 with compounds of the BDM-2 series in the range of 100 to 200. These variations reflect large differences in the dual biochemical activities of these compounds, namely in their ability to inhibit IN–LEDGF/p75 interaction. As shown on Table 4, there is some striking correlation between the IC_50_ value of an INLAI in its ability to disrupt the IN–LEDGF/p75 complex and its ARV potency in SR assay: the INLAI compound with the best IC_50_ in disrupting IN–LEDGF/p75 interaction, MUT871 is also the most potent in ARV activity in SR assay. The weakest compound to inhibit IN–LEDGF/p75, S-I-82 is also the weakest in ARV activity in SR assay.

**Table 4.** Antiretroviral activity of compounds of the BDM-2 series tested by multiple-round (MR) and single-round (SR) infection assays.

| Compound | Biochemistry | NL4-3 infection assays (MT4 cells) |  |  |
| --- | --- | --- | --- | --- |
|  | IN-LEDGF/p75<br>IC<sub>50</sub> (μM) | Single-round<br>EC<sub>50</sub> (μM) | Multiple-round<br>EC<sub>50</sub> (μM) | EC<sub>50</sub> ratio<br>SR/MR |
| **RAL** | NT | 0.0035 ± 0.0010 | 0.014 ± 0.009 | 0.25 |
| **DTG** | NT | 0.00044 ± 0.00013 | 0.0027 ± 0.0013 | 0.16 |
| **BI-224436** | 0.090 ± 0.019 | 1.4 ± 0.3 | 0.051 ± 0.017 | 27 |
| **S-I-82** | 0.82 ± 0.38 | 9.2 ± 1.1 | 0.012 ± 0.003 | 767 |
| **BDM-2** | 0.047 ± 0.014 | 1.4 ± 0.3 | 0.0087 ± 0.0028 | 161 |
| **MUT871** | 0.014 ± 0.0047 | 0.63 ± 0.01 | 0.0031 ± 0.0010 | 203 |
| **MUT872** | 0.17 ± 0.019 | 8.8 ± 0.3 | 0.045 ± 0.012 | 196 |
| **MUT884** | 0.062 ± 0.0094 | 3.9 ± 0.8 | 0.015 ± 0.005 | 260 |
| **MUT916** | 0.046 ± 0.016 | 2.1 ± 0.2 | 0.020 ± 0.009 | 105 |

Determination of the ratio SR versus MR and correlation with IN-LEDGF/p75 interaction IC<sub>50</sub>; comparison with two INSTIs, RAL and DTG, and with two other INLAIs, BI-224436 and S-I-82. Data are mean ± SD of >5 independent experiments. NT, not tested.

### Antiretroviral activity of BDM-2 series on polymorphic recombinant viruses and on primary HIV-1 isolates

The LEDGF-binding pocket, binding site of INLAIs, overlaps with a hot spot of polymorphism on IN amino acids 124 and 125 [29]. Hence it was important to verify that BDM-2 compounds conserved potent ARV activity with all IN 124/125 polymorphisms tested. First, we built 15 different 124/125 polymorphisms in an IN NL4-3 background that covered 98% of all clade polymorphisms described at these positions [32]. Conservation of ARV activity was estimated by the EC_50_ fold change (FC) relative to NL4-3 *TT* at 124/125 positions taken as reference. Results on Table 5A show that compounds of the BDM-2 series conserved their ARV potency on almost all 15 different polymorphisms tested, with FC mostly around 1. For BDM-2, FC was 1 for all polymorphisms expect for *AV* with FC of 2. For MUT871, only the 124/125 polymorphisms *GT*, *GA* and *NA* showed FC of 2, 3 and 2 respectively. For MUT916, the only polymorphism with FC >1 was *NA* with FC of 4. FC changes of 3-fold or less are considered modest in nature, demonstrating that this BDM-2 series is associated with broad polymorph coverage. In contrast S-I-82 lost most of its ARV activity when an *N* residue is found at position 124, with a FC of 10 (meaning conservation of 10% activity only) for *NA* at 124/125 residues, and a FC of 5 for *NT* or *NV* at 124/125 positions in the IN sequence and a FC of 4 for *SA* 124/125 polymorphism.

**Table 5.** ARV activity of compounds of the BDM-2 series against hot spot 124-125 polymorphism.

A. Recombinant polymorphic viruses on the hot spot 124-125 polymorphism
| Compound | EC<sub>50</sub> FC of MR infection assays in MT4 cells with position 124-125 polymorphic viruses vs. NL4-3 TT |  |  |  |  |  |  |  |  |  |  |  |  |  |
| --- | --- | --- | --- | --- | --- | --- | --- | --- | --- | --- | --- | --- | --- | --- |
|  | AA | AM | AT | AV | GT | GA | NA | NT | NV | SA | ST | TA | TM | TV |
| **DTG** | 1 | 2 | 1 | 2 | 2 | 2 | 1 | 1 | 2 | 1 | 2 | 1 | 2 | 2 |
| **BI-224436** | 1 | 0.4 | 1 | 1 | 2 | 1 | 1 | 1 | 1 | 1 | 1 | 1 | 0.4 | 1 |
| **S-I-82** | 1 | 1 | 1 | 2 | 3 | 3 | 10 | 5 | 5 | 4 | 2 | 2 | 1 | 1 |
| **BDM-2** | 1 | 0.4 | 1 | 2 | 1 | 1 | 1 | 1 | 1 | 1 | 1 | 1 | 0.3 | 1 |
| **MUT871** | 1 | 0.4 | 1 | 1 | 2 | 3 | 2 | 1 | 1 | 1 | 1 | 1 | 0.3 | 1 |
| **MUT872** | 1 | 0.3 | 1 | 0.5 | 2 | 2 | 1 | 1 | 1 | 1 | 1 | 1 | 0.2 | 1 |
| **MUT884** | 0.4 | 0.2 | 1 | 0.5 | 2 | 1 | 2 | 1 | 1 | 1 | 1 | 1 | 0.2 | 1 |
| **MUT916** | 1 | 0.2 | 0.5 | 0.4 | 1 | 1 | 4 | 1 | 0.4 | 1 | 1 | 1 | 0.2 | 0.4 |

B. NL4-3 with integrase from HIV-1 primary isolates (hot spot 124-125 polymorphism)
| Compound | EC<sub>50</sub> FC of MR infection assays in MT4 cells with integrases from primary isolates vs. NL4-3 |  |  |  |  |  |  |  |  |  |  |  |  |  |  |  |  |
| --- | --- | --- | --- | --- | --- | --- | --- | --- | --- | --- | --- | --- | --- | --- | --- | --- | --- |
|  | AA |  |  |  |  |  | GA |  | NA |  |  |  | NV | SA |  | TA |  |
|  | KER2008<br>(A) | TZA125<br>(C) | E0836M4<br>(D) | NP1525<br>(CRF01) | NP1695<br>(CRF02) | CAM2BB<br>Y (CRF02) | KSM4030<br>(A) | vGA<br>(C) | NP1538<br>(B) | 99ET14<br>(C) | vNA_1<br>(B) | vNA_2<br>(CRF02) | vNV<br>(B) | vSA_1<br>(B) | vSA_2<br>(CRF02) | vTA_1<br>(B) | vTA_2<br>(CRF02) |
| **DTG** | 1 | 2 | 2 | 2 | 2 | 1 | 2 | 2 | 1 | 2 | 2 | 2 | 1 | 1 | 2 | 1 | 1 |
| **BI-224436** | 1 | 0.5 | 1 | 1 | 2 | 2 | 1 | 2 | 1 | 1 | 2 | 2 | 1 | 3 | 2 | 2 | 2 |
| **S-I-82** | 1 | 1 | 1 | 2 | 1 | 2 | 6 | 4 | 1 | 210 | 23 | 23 | 34 | 12 | 4 | 3 | 3 |
| **BDM-2** | 1 | 0.4 | 1 | 1 | 2 | 2 | 2 | 4 | 2 | 1 | 2 | 3 | 2 | 3 | 3 | 2 | 1 |
| **MUT871** | 1 | 1 | 2 | 1 | 3 | 2 | 2 | 5 | 2 | 2 | 3 | 3 | 3 | 4 | 3 | 2 | 2 |
| **MUT872** | 1 | 0.4 | 1 | 1 | 1 | 1 | 2 | 4 | 2 | 1 | 1 | 2 | 1 | 2 | 2 | 2 | 2 |
| **MUT884** | 0.5 | 0.3 | 1 | 1 | 2 | 1 | 1 | 5 | 3 | 1 | 3 | 4 | 2 | 2 | 2 | 1 | 2 |
| **MUT916** | 1 | 0.4 | 1 | 1 | 2 | 1 | 1 | 5 | 2 | 2 | 4 | 10 | 1 | 3 | 4 | 2 | 2 |

Antiretroviral activity against (A) recombinant polymorphic viruses constructed in NL4-3 background and based on the hot spot polymorphism of IN 124/125 residues, and (B) against primary HIV-1 isolates. Evaluation of conserved activity by EC<sub>50</sub> fold change (FC) determination with respect to wt NL4-3 virus. FC >10 are highlighted in dark grey; FC between 5 and 10 in light grey. The different polymorphisms on 124/125 residues of the primary HIV isolates are indicated, as well as the clade of each of the HIV-1 primary isolates indicated between brackets. Comparison with one INSTI, DTG and two other INLAI, BI-224436 and S-I-82 used as controls. Data are mean ± SD of >5 independent experiments.

The search about conservation of ARV potency of BDM-2 compounds on polymorphic viruses was completed by analyzing ARV activity of BDM-2 compounds on several primary isolates exhibiting different polymorphisms at positions 124/125. As shown on Table 5B, BDM-2 compounds conserved excellent ARV activity on all 14 primary isolates tested, with FC of 1 to 3 most frequently except for the primary isolate vGA for all BDM-2 compounds with FC of 4 or 5 and for MUT916 with FC of 10 for the primary isolates vNA_2. In contrast, S-I-82 had important losses, particularly in all primary isolates with *NA* or *NV* in positions 124/125 with FC of 210, 23, 23, 14 respectively and a FC of 12 for the primary isolate vSA_1.

### BDM-2 is fully active on viruses resistant to all classes of current drugs

Importantly we checked that BDM-2 as representative lead compound of this series was fully active on all resistant viruses to drugs currently used in clinic. To do so we used both recombinant resistant viruses constructed by introducing described resistance mutations in an NL4-3 background and resistant clinical isolates. As shown on Table 6, this was indeed the case on recombinant and primary clinical isolates resistant to INSTIs, NRTIs, NNRTIs and PIs with a FC of 1 for all resistant viruses except FC of 2 for one Reverse Transcriptase inhibitor resistant virus. The studied mutations had no significant effect on other compounds of the BDM-2 series as well.

**Table 6.**
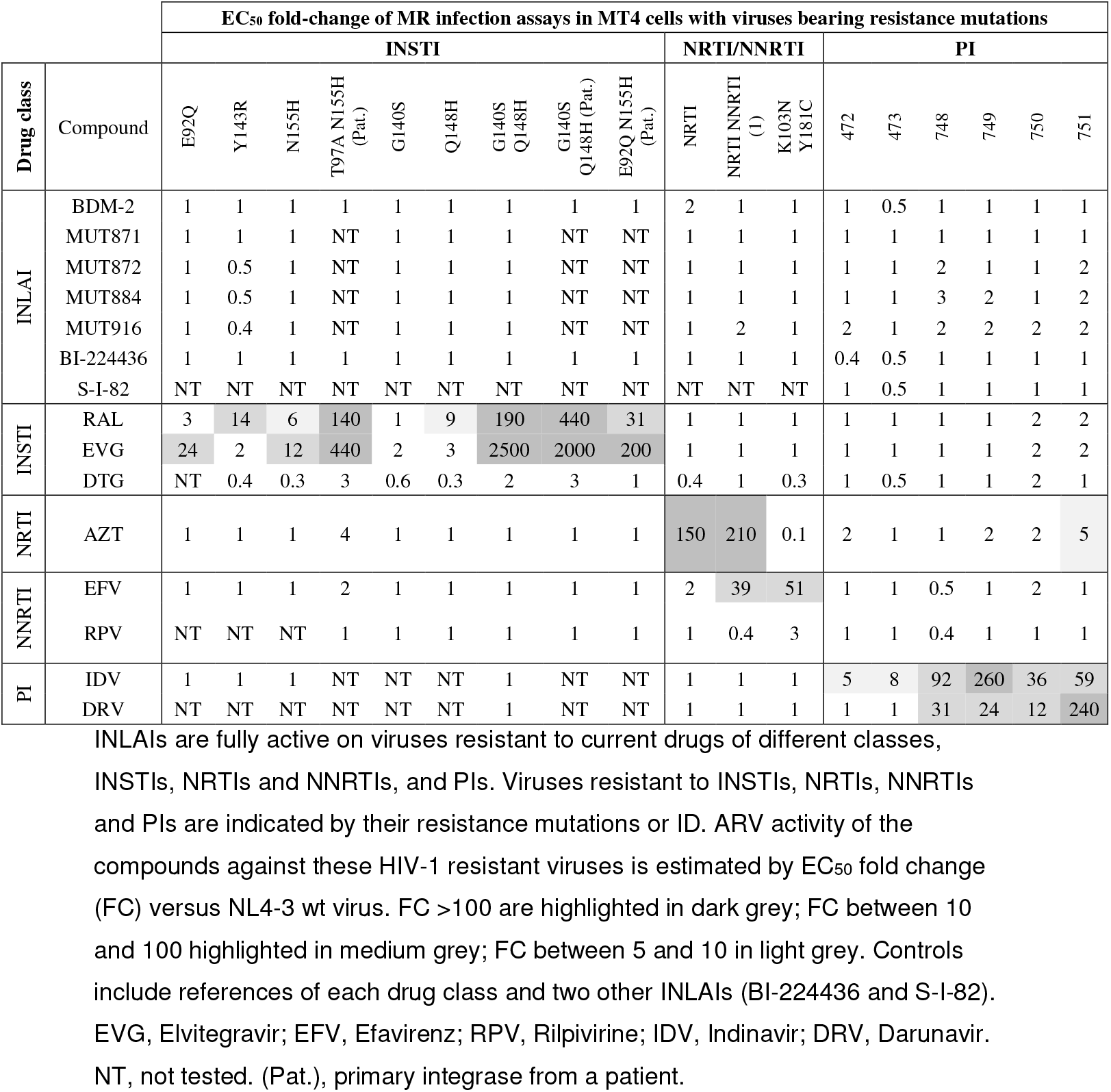
Antiretroviral activity (ARV) of compounds of the BDM-2 series against viruses resistant to other drug classes.

### Structure of BDM-2 bound to IN-CCD

The overall structures of unliganded (PDB 4LH4 [29] and liganded IN-CCD are very similar (see Fig 2), including the general topology of the LEDGF/p75 binding site (Fig 2A). BDM-2 is unambiguously defined in the density (Fig 2B). BDM-2 is anchored in the IN-CCD pocket through H-bonds with Glu170, His171 main chain and Thr170 side chain (Fig 2C). BDM-2 is buried in a hydrophobic pocket defined by residues Leu102, Thr124, Thr125, Ala128, Ala129, Trp132, Ala169, Gln168 and Met178 (Fig 2C). In the unliganded IN-CCD [29], the LEDGF/p75 binding pocket is filled with water molecules (Fig 2D). The superposition of the LEDGF binding pocket in the unliganded (Fig 2D) and liganded form (Fig 2E) shows a large movement of Glu170 side chain upon BDM-2 binding (Fig 2F).

**Fig 2.**
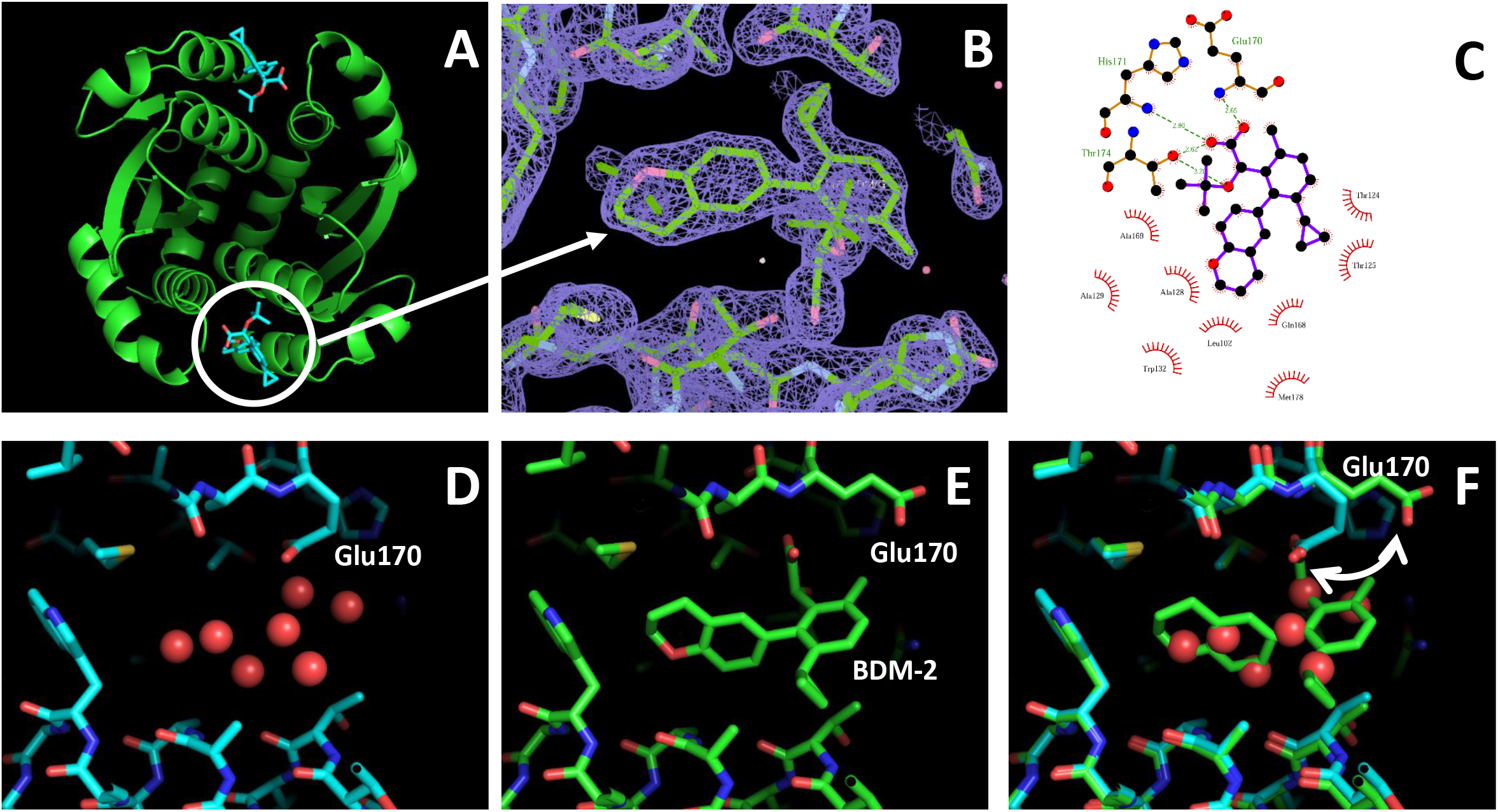
BDM-2 co-crystallized with IN-CCD. Comparison of the IN-CCD structures in absence (4LH4) and in presence of BDM-2. (A) Full view of the IN-CCD dimer, Main chain in green and BDM-2 in blue. (B) Enlargement of the BDM-2 binding pocket showing the 2Fo-Fc electronic density map. (C) BDM-2 – IN-CCD specific interactions. (D) Close view of the structure of the IN-CCD LEDGF binding pocket without ligand (PDB 4LH4 [29]). The pocket is filled with water molecules (red spheres). (E) Close view of the structure of the IN-CCD LEDGF binding pocket with BDM-2. (F) superposition of the ligand binding pocket without and with BDM-2. The arrow shows the movement of Glu170 side chain upon BDM-2 binding.

### HIV-1 mutants selected for resistance to INLAI compounds of BDM-2 series

Selection of *in vitro* resistance to BDM-2, as lead compound representative of the BDM-2 series, was based on dose-escalation method and evaluated in parallel with BI224436, another INLAI based on different chemical scaffold, as well as with RAL, and Nevirapine (NVP) as reference antiretroviral compounds, under the same conditions. The progress of virus replication in the presence of stepwise increasing concentrations of inhibitors was monitored over time (Fig 3). The kinetics of BDM-2 resistance selection, as well as that of BI-224436, was similar to that of NVP and slightly faster than RAL. These results confirmed that BDM-2 and more generally INLAIs display low genetic barrier to resistance, similarly to NVP and possibly lower than RAL.

**Fig 3.**
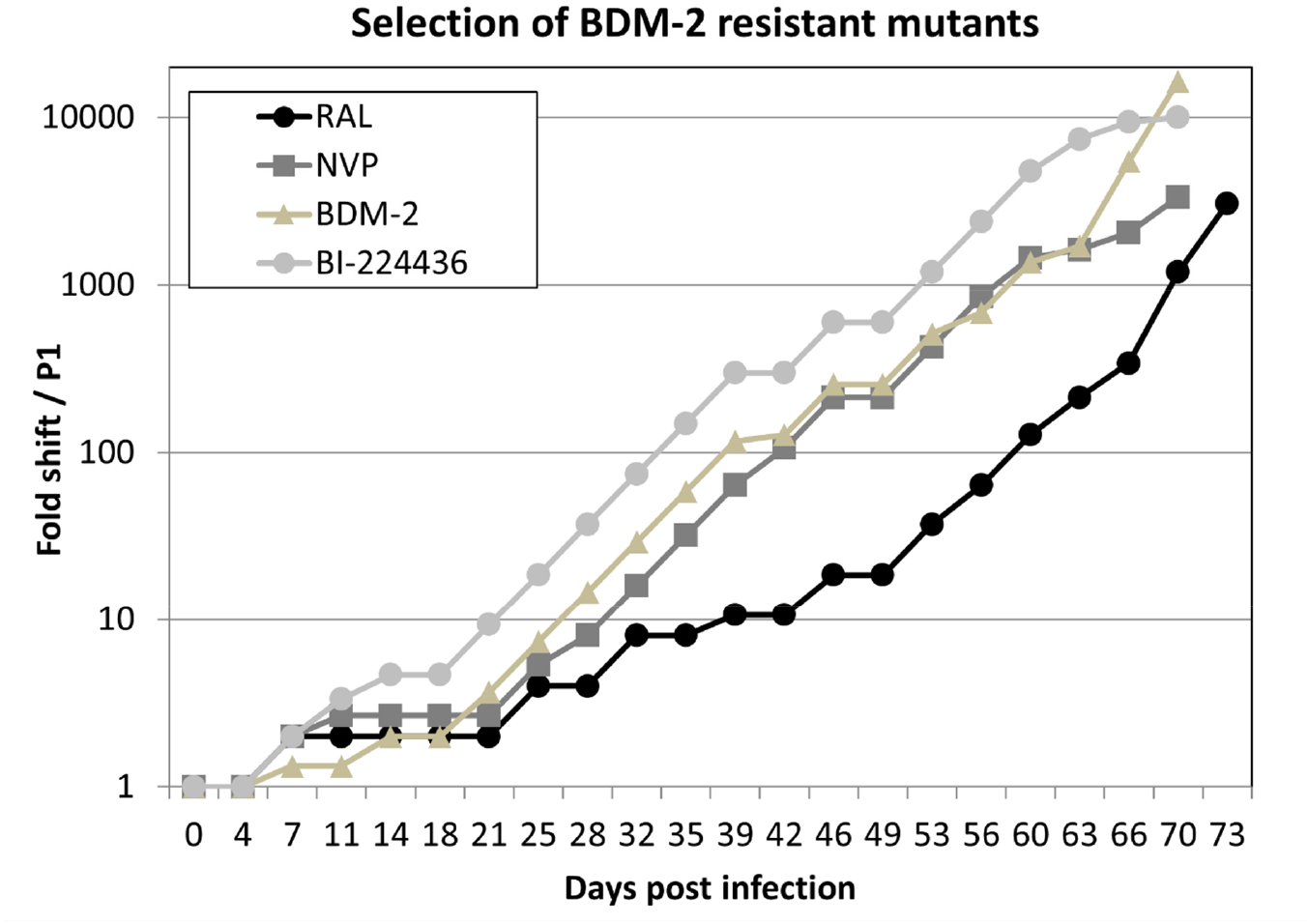
Selection of BDM-2 resistant mutants. Selection of resistant mutants to BDM-2 and BI-224436 in MT4 cells infected with HIV-1 NL4-3. This selection was based on dose-escalation drug method and evaluated in parallel with RAL, and Nevirapine (NVP) as reference antiretroviral compounds, under the same conditions. The resistance selection process was initiated at the EC_50_ concentration for each drug. The progress of virus replication in the presence of stepwise increasing concentrations of inhibitors was monitored over time during 73 days by fold change in EC_50_ of all drugs used in these experiments.

The IN-coding sequences have been analyzed in selected HIV-1 resistant viruses isolated at various times of the resistance selection process (Table 7). During the selection for BDM-2 resistant mutants, the occurrence frequency of the different mutations was estimated as a percentage of the total number of mutations detected on IN sequences as a bulk. Early mutations detected at day 18 were mutations close to the INLAI binding site in the IN-CCD: A128T, the most frequent one found in 50% of IN sequences, and Y99H found in 36% of IN sequences. Interestingly a low level (7%) of N222K mutation in the CTD domain was also detected. At intermediate selection time, day 42, this landscape of INLAI resistance mutations did not change much. The most detrimental mutations such as T174I, and to a lesser degree A129T, occurred late during the selection process, between days 66 and 73, while previously detected mutations A128T and N222K were still present in IN sequences. Identical mutations with similar kinetics were also detected during the selection process with BI-224436 (see Fig 3 and Table 7).

**Table 7.** Kinetics of occurrence of INLAI resistant mutants.

| **A** | Kinetic (average %) |  |  |
| --- | --- | --- | --- |
|  | Early | Intermediate | Final |
| Y99H | 36 | 31 | 23 |
| T125A | 0 | 0 | 3 |
| A128T | 57 | 50 | 30 |
| A129T | 0 | 0 | 10 |
| E170G | 0 | 0 | 3 |
| H171Q | 0 | 0 | 3 |
| L172I/F | 0 | 0 | 3 |
| T174I | 0 | 0 | 10 |
| N222K | 7 | 19 | 13 |

| **B** | Kinetic (average %) |  |  |
| --- | --- | --- | --- |
|  | Early | Intermediate | Final |
| Y99H | 0 | 11 | 13 |
| A128T | 40 | 33 | 38 |
| E170G | 20 | 11 | 0 |
| H171Q | 20 | 11 | 13 |
| T174I | 0 | 22 | 25 |
| N222K | 20 | 11 | 13 |

Occurrence frequency of the different resistant mutations detected in IN sequence during the selection process with (A) BDM-2 and (B) BI-224436 were identified at three different time periods, early at 18 days, intermediate at 42 days and final occurrence between 66 to 73 days. The frequency of each identified mutation was estimated as a percentage of the total number of mutations detected on IN sequences as a bulk.

Major single point mutations identified during the resistance selection process to BDM-2 were introduced in an HIV-1 NL4-3 background and the FC in the EC_50_ of each resistant mutant relative to wild-type virus was determined (see Table 8). The most detrimental mutation for BDM-2 compounds and all the other INLAIs investigated was T174I with fold change of 210 for BDM-2, 544 for BI-224436 and 15 for S-I-82. All the other single mutations detected during the selection process, Y99H, A128T, H171Q, N222K yielded much weaker resistance with FC of 4, 4, 3 and 2 respectively. All INLAIs investigated shared similar resistance profile to that of compounds of the BDM-2 series. These results are consistent with resistance profile for previously reported INLAIs. Interestingly, Elvitegravir (EVG) as INSTI representative of current drugs used in clinic, conserved full activity on all INLAI resistant mutants.

**Table 8.** Antiretroviral activity of compounds of the BDM-2 series against some frequent INLAI resistance single point mutants selected by serial passages.

| Compound | EC<sub>50</sub> (nM)<br>NL4-3 | EC<sub>50</sub> fold-change of NL4-3 with resistance mutations vs. wt |  |  |  |  |
| --- | --- | --- | --- | --- | --- | --- |
|  |  | A128T | Y99H | H171Q | T174I | N222K |
| EVG | 2.2 ± 1.2 | 1 | 1 | 1 | 1 | 1 |
| BI-224436 | 51 ± 17 | 6 | 2 | 4 | 706 | 2 |
| S-I-82 | 12 ± 3 | 4 | 2 | 2 | 15 | 1 |
| BDM-2 | 8.7 ± 2.8 | 4 | 4 | 3 | 253 | 2 |
| MUT871 | 3.1 ± 1.0 | 2 | 4 | 6 | 323 | 3 |
| MUT872 | 45 ± 12 | 7 | 2 | 1 | 15 | 2 |
| MUT884 | 15 ± 5 | 5 | 2 | 1 | 12 | 3 |
| MUT916 | 20 ± 9 | 4 | 3 | 1 | 14 | 3 |

ARV activity is estimated by Fold Change (FC) versus NL4-3 wt virus; FC >10 are highlighted in dark grey; FC between 5 and 10 in light grey. comparison with one INSTI, EVG, and two other INLAIs used as references (BI-224436 and S-I-82).

Most of the INLAI resistance mutations were found as expected in the IN-CCD domain in or close to the LEDGF binding pocket, binding site of all INLAIs. However, one mutation repetitively found during the selection mutation process, N222K, was found in the IN-CTD, which underlined the importance of chemical interactions interference between INLAI molecules and some amino acids of the IN-CTD domain.

### Replication capacity of INLAI resistant viruses

We then analyzed the impact of the various INLAI resistance mutations on the replication capacity of mutated viruses. As shown in S1 Fig, the most detrimental mutation T174I had also the greater negative impact on replication capacity that was estimated in average to 29% of that of wild type virus. All the other mutations that resulted in moderate resistance, Y99H, N222K, A128T and H171Q, had also lower impact on replication capacity with 90%, 70%, 70%, 70% with regard to wild type virus respectively.

### No antagonism of BDM-2 with different ARV drugs used in clinic

All efficient treatments against HIV-1 infection are combination therapies of several (three usually for the first generation of Highly Active Anti-Retroviral Therapy (HAART)) ARV drugs with different mode of action. It was thus important, in the putative perspective of using BDM-2 the lead compound of this series as a potentially novel component of such combination therapies, to determine if BDM-2, did not antagonize other ARV drugs. Combination studies of BDM-2 with approved ARV drugs were performed in MT4 cells infected by HIV-1 NL4-3. In Fig 4A and B are shown examples of MacSynergy plots obtained after combination of BDM-2 with the protease inhibitor Lopinavir (LPV) (A) or the INSTI Elvitegravir (B). These plots indicated that there was no antagonism of ascending concentration of BDM-2 with the ARV activity of increased concentration of Lopinavir or Elvitegravir and vice versa. In Fig 4C this type of analysis was extended to a panel of 16 FDA-approved drugs from the ARV pipeline (3 PIs, 6 NRTIs, 4 NNRTIs and 3 INSTIs). Compared to the synergy or antagonism obtained by combinations Ribavirin + DDI (synergy) or Ribavirin + AZT (antagonism used as controls (Fig 4D), BDM-2 displayed no antagonism with none of the 16 drugs investigated, relatively strong synergy with Lopinavir and moderate synergy for some of these 16 drugs including the INSTIs, EVG, RAL or DTG (Fig 4C).

**Fig 4.**
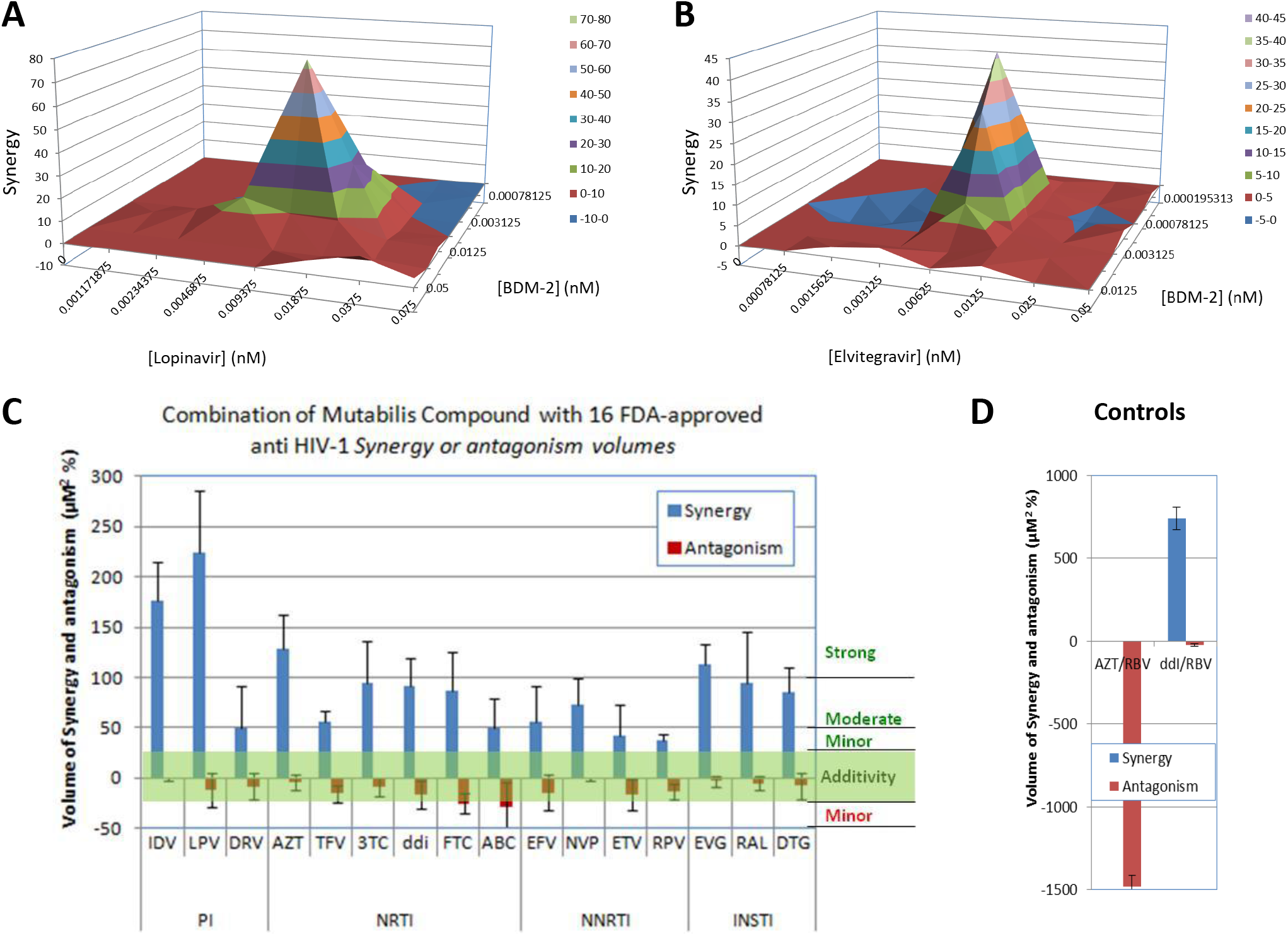
Study of various combinations of BDM-2 with 16 different approved ARV drugs by MacSynergy plot to analyze synergy or antagonism between BDM-2 and these drugs. A & B. MacSynergy plots of ARV activity obtained after treatment of NL4-3-infected MT4 cells by combination of drugs, (A) with BDM-2 + Lopinavir, (B) with BDM-2 + Elvitegravir, (C) quantitative results of synergy or antagonism volume by combination of BDM-2 with 16 FDA-approved drugs of different classes, (D) volume of synergy or antagonism of combinations of ddI + Ribavirine (RBV) (synergy) and AZT + RBV (antagonism) used as controls.

## Discussion

In this report, we described the biochemical and virological properties of INLAI compounds of the BDM-2 series (BDM-2s), novel and potent inhibitors of HIV-1 replication. BDM-2s belong to a novel class of ARVs of the INLAI family recently developed, but not yet in clinical use. INLAIs are allosteric inhibitors of HIV-1 IN that inhibit IN-LEDGF/p75 interaction. ARV activity of BDM-2s is more powerful than that of RAL and is about the same magnitude as DTG the best ARV drug currently in clinical use. BDM-2s are among the most potent INLAIs ever described. They have potent ARV activity in both T-lymphoblastoid cell lines and primary human T-lymphocytes. Moreover, they display consistent ARV activity across most of HIV-1 polymorphisms at the hotspot 124/125 IN residues and on most HIV-1 clinical isolates of various subtypes tested. Like NNRTIs and in contrast to INSTIs, all compounds of this INLAI family of inhibitors are devoid of significant activity on HIV-2 or SIV viruses. BDM-2s conserved full activity against a panel representative of mutated viruses resistant to other classes of FDA-approved drugs used in clinic. The selectivity ratio (CC_50_/EC_50_) of BDM-2s was very high in several cell types showing low cytotoxicity. Importantly, no antagonism was found between BDM-2s and the clinically approved ARV drugs, and a somewhat strong or moderate synergy was found with the PI Lopinavir and INSTIs respectively. Mutants resistant to BDM-2s were selected by dose escalation method and several point mutations were identified, T174I being the most detrimental with a high fold shift in EC_50_ compared to wild type viruses. All INLAIs described up to now were found sensitive to these resistance mutations at various extent, with similarly T174I being the most detrimental mutation. This indicates that the genetic barrier to resistance of INLAIs described to date is quite low, and this is an important point to be improved if one wants to use this class of drugs in combination with other ARV classes. With improved barrier of resistance, this drug class could be efficiently used in combination with all the other classes of ARV drugs that retain full activity on BDM-2 resistant mutants (not shown). INLAI compounds with improved genetic barrier to resistance and BDM-2s in particular, could be of great interest for HIV-1 infected patients that are in therapeutic failure because of multi-resistance to current ARV drugs.

Mutation N222K was the only INLAI resistance mutation found in the IN CTD, outside the INLAI binding pocket located in the CCD. Interestingly, M. Kanja *et al*. identified, in the IN CTD, a new functional motif constituted by four noncontiguous amino acids (N_222_K_240_N_254_K_273_) [33], including this position N222. This NKNK motif strictly conserved in all natural sequences of HIV-1 group M, generates a positive surface potential that is important for the nuclear import of reverse-transcribed genomes. This motif could also be involved in functions of integrase different of integration, such as reverse transcription. It is located in a region of the CTD in which IN interacts with Reverse Transcriptase [34,35]. The INLAI resistance mutation we found at this position, N222K, seems to slightly increase integration [33]. It is tempting to speculate that this effect might be related to the mechanism generating weak but consistent resistance to INLAIs.

Co-crystallization of BDM-2 compounds with the IN CCD domain as successfully achieved here, led us to elucidate how BDM-2 and other compounds of this series bind within their INLAI binding site in the CCD.

INLAI compounds, besides their interest as a new class of ARV drug with a novel mechanism of action, attract also great interest for their ability to dissociate the complex IN-LEDGF/p75. Indeed, LEDGF/p75 is the main cellular cofactor of IN that target integration of HIV-1 in actively transcribed genes (for review see [36]). Zeger Debyser’s group postulated that inhibitors of IN-LEDGF interaction might be useful in a block-and-lock strategy [37,38] by inhibiting viral integration at actively transcribed genes and retargeting HIV-1 integration to sites that are less susceptible to reactivation. However, we should take into account that all INLAIs display a double mechanism of action, inhibition of HIV-1 integration by disrupting IN-LEDGF/p75 interaction and inhibition of maturation of viral particles at post-integration step by enhancing aberrant IN multimerization. This later activity of enhancement of IN aberrant multimerization at post-integration step is much more potent than that at integration based on the disruption of the complex between IN and LEDGF/p75. This is only this later activity of IN-LEDGF/p75 complex dissociation that could be useful for a block-and-lock strategy. Consequently, only INLAIs that could have sufficiently potent activity in disruption of the complex between IN and LEDGF/p75 could be considered for such block-and-lock strategy. Toward this goal it is essential to evaluate by single round infection assay the INLAIs that could be selected for such a strategy. From this point of view, MUT871 and BDM-2 can be considered as good candidates. MUT871 dissociates the IN-LEDGF/p75 complex with IC_50_ of 14 nM and has ARV activity in single round assay with EC_50_ of 630 nM. BDM-2 dissociates the IN-LEDGF/p75 complex with IC_50_ of 47 nM and has ARV activity in single round assay with EC_50_ of 1.4 μM.

To date there is still no INLAI in advanced clinical investigation for ARV proof of concept studies in patients infected with HIV-1 in phase IIa and phase III clinical trials. BI-224436 from Boehringer Ingelheim originally was introduced in phase I clinical trial, but this trial was interrupted for unknown reason [39], presumably because of toxicity issues. Gilead reported at a CROI meeting in 2017 [31] a very active INLAI compound with a benzothiazole core group, GS-9822, however this compound provoked renal and urinary bladder toxicity which precluded further clinical investigation. VIIV also developed a new series of very active INLAIs, tetrahydronaphtyridines [40], however these compounds have not yet reached clinical investigation. ST Pharm (Seoul, South Korea) reported also a very active INLAI compound, Pirmitegravir or STP0404 [15], and recently presented the results of a clinical phase I study at the 24th AIDS Conference [41]. Well tolerated and with favorable pharmacokinetics suitable for once-daily low dose regimen, the compound is expected to move on to a phase IIa trial in Q4, 2022.

After completion of preclinical studies (data not shown), BDM-2 has been further investigated in a single ascending dose phase I clinical trial investigating Safety, Tolerability and Pharmacokinetics in healthy male subjects completed as soon as 2020. The results of this trial were reported June 17^th^ 2020 in clinicaltrials.gov without any serious adverse event and few mild adverse events ([42] and manuscript in preparation). Therefore, BDM-2 is together with Pirmitegravir the most advanced INLAI in investigation today in man, which supports further clinical investigation.

## Materials and methods

### Compound synthesis

BDM-2, MUT871, MUT872, MUT884 and MUT916 were synthesized at Biodim as described in patent application WO2015/001125A1, according to examples 13, 38, 24, 40 and 44 respectively. Reference BI-224436 was prepared as described in patent application WO2009/062285A1, according to compound 1144; S-I-82 was prepared as described in WO2013/062028A1 according to compound I-82.

### Reference compounds

Control compounds such as Nevirapine (NVP), Efavirenz (EFV) and AZT were obtained from the NIH AIDS Research and Reference Reagent Program. Raltegravir (RAL) Dolutegravir (DTG), Nevirapine (NVP), Indinavir (IDV), AZT, Ribavirin (RBV), Lopinavir (LPV), Darunavir (DRV), Tenofovir (TFV), Lamivudine (3TC), Didanosine (ddI), Emtricitabine (FTC), Abacavir (ABC), Efavirenz (EFV), Etravirine (ETV), Rilpivirine (RPV) and Elvitegravir (EVG) were purchased from Selleck Chemicals.

### Molecular biology and biochemistry

Epitope-tagged proteins used in IN–LEDGF/p75, IN-CCD–LEDGF-IBD interactions and in IN multimerization assays were constructed and purified as described previously [8].

HTRF^®^-based CCD–IBD interaction, IN–LEDGF/p75 interaction and IN multimerization assays were performed as described previously [8].

### Virology

#### Cell culture

MT4 cells were obtained through the AIDS Research and Reference Reagent Program, Division of AIDS, NIAID, NIH. MT4 cells were grown in RPMI 1640 supplemented with 10% heat-inactivated fetal calf serum and 100 IU/mL penicillin, and 100 μg/mL streptomycin (Invitrogen) to obtain RPMI-complete medium.

#### Virus strains and recombinant HIV-1 molecular clones

HIV-1 NL4-3 and NL4-3Δ*env*-luc molecular clones were obtained from the NIH AIDS Research and Reference Reagent Program.

#### Viral stocks

Were prepared and quantified as described previously [8] in 293T cells, single-round viral stocks were produced by co-transfecting pNL4-3Δ*env* with VSV-G envelope expression vector [8]. HIV-1 resistant mutants were constructed as described previously [29].

#### Antiviral assay in MT4 cells

MT4 cells growing exponentially at the density of 10^6^/mL were infected with HIV-1 strain NL4-3 at a MOI (multiplicity of infection) of 0.001 for 2 h in the presence of different concentrations of compounds, and the CellTiter-Glo^®^ luminescent reagent (Promega) was used to quantify cell viability as described previously [8]. In order to evaluate the effect of the human serum on the ARV potency on INLAIs and other ARV compounds used as reference, we measured their EC_50_ and EC_90_ from the dose response curve in the presence of various concentrations of human serum between 0 to 50% and linearly extrapolated their value at 100% human serum for PA-EC_90_ determination.

#### Replication-defective-HIV assay (single round infection assay)

MT4 cells (growing exponentially at the density of 10^6^/mL) were infected with VSV-G-pseudotyped NL4-3Δ*env*-luc at a MOI of 0.0001 and Luciferase expression was quantified after two days using the One-Glo™ luciferase assay (Promega) as described previously [8].

#### Cytotoxicity assays

Growth inhibition was monitored in a proliferating human T-cell line (MT4) with different concentrations of compounds, using the CellTiter-Glo^®^ luminescent reagent (Promega) as previously described [8].

#### Resistant virus selection

MT4 cells infected with HIV NL4-3 isolate were cultured in the presence of BDM-2 at the EC_50_ concentration determined earlier. At each passage, cells from original culture in the presence of inhibitor were mixed with equal amount of no-drug control cells to propagate, and viral replication was monitored by the production of p24 antigen in the supernatant. Inhibitor concentration was gradually increased at each passage. At three different times, early (day 18), intermediate (day 42) and final passage (day 73), viral RNA was extracted using QIAexpress (Qiagen) and IN sequences were determined by RT-PCR as a bulk. The quantitative estimate of each mutation induced at each passage mentioned above was determined as a percentage of total mutations detected in the IN sequences as a bulk.

#### Viral replication capacity

HIV-1 recombinant viruses harboring various single point INLAI resistant mutations (A128T, Y99H, N222K, T174I, H171T, L102F and T124D) in an NL4-3 background were used to infect MT4 cells with equivalent p24 quantity, and compared with infection of MT4 cells in same conditions with identical p24 quantity of wild type NL4-3 and of some INSTI resistant viruses (G140S and Q148H). Replication kinetics of the INLAI resistant variants were compared and viral production was determined by p24 assay daily for 5 days.

#### Combination antiviral activity assays

BDM-2 as lead compound representative of the series was used in these experiments. Combination studies were performed in MT4 cells infected with NL4-3, as previously described [2]. Multiple concentrations of BDM-2 were tested in checkerboard pattern in the presence and absence of dilutions of 16 representative approved anti-HIV drugs of different classes. Compound combinations were analyzed by calculations to quantify deviation from additivity at the 50% level. Data were analyzed as described by Prichard and Shipman by using the MacSynergy II program [43]. Synergy volumes in the range of - 25 to + 25 define additivity; −50 minor antagonism, + 50 minor synergy, and up to 100 and > 100 moderate and strong synergy respectively. Combinations of Ribavirine (RBV) with AZT or DDI were used as controls for antagonism and synergy respectively.

#### Structural studies with IN-CCD

Expression and purification of the HIV-1 IN CCD F185K (50-212) were performed as previously described [8] and used for crystallization.

Briefly, the sequence of HIV IN CCD F185K (50-212) with a N-terminal HVR3C Prescission protease cleavage site was cloned in pGEX-6P by enzymatic restriction through BamHI and XhoI sites and overexpressed in E. coli BL21(DE3) STAR cells. 4 L of LB (100 mg/L ampicillin) were induced at an OD600 of 0.6-0.8 with 0.5 mM IPTG for 18 hours at 18ºC. Cells were harvested, resuspended in lysis buffer (50 mM HEPES pH 7.5, 0.5 M NaCl, 5 mM MgCl2, 5 mM DTT) at a ratio of 10 mL of buffer per gram of biomass, in presence of 1 mM PMSF and Roche Complete inhibitor cocktail tablets to avoid protease degradation. Lysis was performed by pulse sonication every 2 s at 40% amplitude for 1 min/g of cells at 4°C. The lysate was then clarified by ultracentrifugation for 1 h at 125,000 g at 4°C. After filtration on cellulose filter 5μ, sample was loaded on GSTrap FF 5 mL column (Cytiva). Column was washed with lysis buffer and proteins were eluted through a cleavage of the GST tag on column with 2 mg of HVR3C prescission protease by incubation for 18 h at 4°C. The sample was then concentrated on Amicon (10kDa MWCO) and further purified using a Superdex 75 10-300 GL column (Cytivia) equilibrated with GF buffer (50 mM MES pH 5.5, 50 mM NaCl, 5 mM DTT). Fractions containing the purest complex (checked by SDS-PAGE) were then pull together and concentrated on Amicon (10 kDa MWCO) and used for crystallization.

Crystals were grown at 20°C by vapor diffusion using 3 μL of protein at 5 mg/mL in 50 mM MES pH5.5, 50 mM NaCl, 5 mM DTT mixed to 3 μL of reservoir solution containing 0.1 M sodium cacodylate pH 6.5, 1.26 M ammonium sulfate, with 500 μL of reservoir solution in the well [29]. Less than a week was needed to get crystals. Then, crystals were soaked by adding 0.1 μL of 12 mM ligand/BDM-2 at 20°C to the drop 8 hours before data acquisition.

After fishing, crystals were cryo-protected in oil (FOMBLIN Y LVAC 14/6 from Aldrich) for a few seconds and cryo-cooled in a stream of liquid nitrogen at −173 °C. Data collection has occurred in house on a Rigaku FR-X rotating anode equipped with a Dectris Eiger R 4M detector. X-ray diffraction images were indexed and scaled with XDS [44]. Structure solving, and refinement were performed using PHENIX program suite [45].

The structure was solved by molecular replacement using Phaser [46]. It was refined with Refine [47]. The ligand BDM2 coordinates and restrains were generated with Elbow [48] and was placed in the structure using LigandFit [49]. Structure superpositions were performed in Coot [50]. All structure drawings were performed with PyMOL [51] and Coot [50]. 2D view of ligand interactions have been generated with LigPlot [52]. Statistics of data scaling and structure refinement are summarized in S2 Table. Structures and structure factors have been deposited in the PDB database with codes 8BV2.

## Acknowledgements

We thank Wandrille Ract-Madoux for support; Ibrahima Guillard for administrative assistance; Juliette Nguyen, Roxane Beauvoir, and Elodie Drocourt for assistance in virology; Jean-Michel Bruneau for assistance in biochemistry; Isabelle Mallet, Nicolas Levy for help in crystallographic data collection and the NIH HIV Reagent Program, Division of AIDS, NIAID, for HIV-1-drug resistant viruses and HIV-1 primary isolates.

The structural part of this work of the Interdisciplinary Thematic Institute IMCBio, as part of the ITI 2021-2028 program of the University of Strasbourg, CNRS and Inserm, was supported by IdEx Unistra (ANR-10-IDEX-0002), and by SFRI-STRAT’US project (ANR 20-SFRI-0012) and EUR IMCBio (ANR-17-EURE-0023) under the framework of the French Investments for the Future Program. We acknowledge the French Infrastructure for Integrated Structural Biology (FRISBI) ANR-10-INSB-0005 and Instruct-ERIC.

## Author contributions

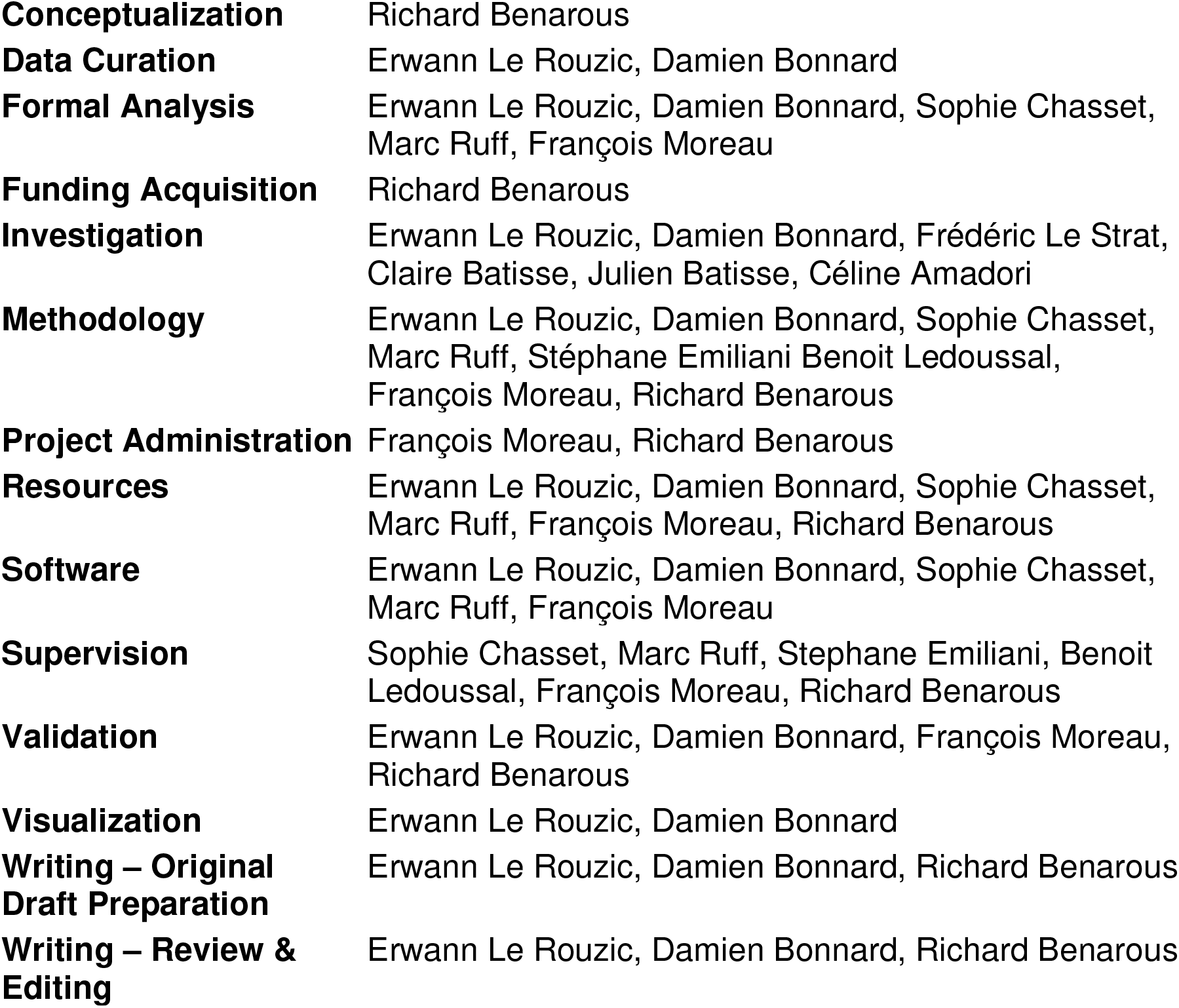

---

**S1 Fig.**
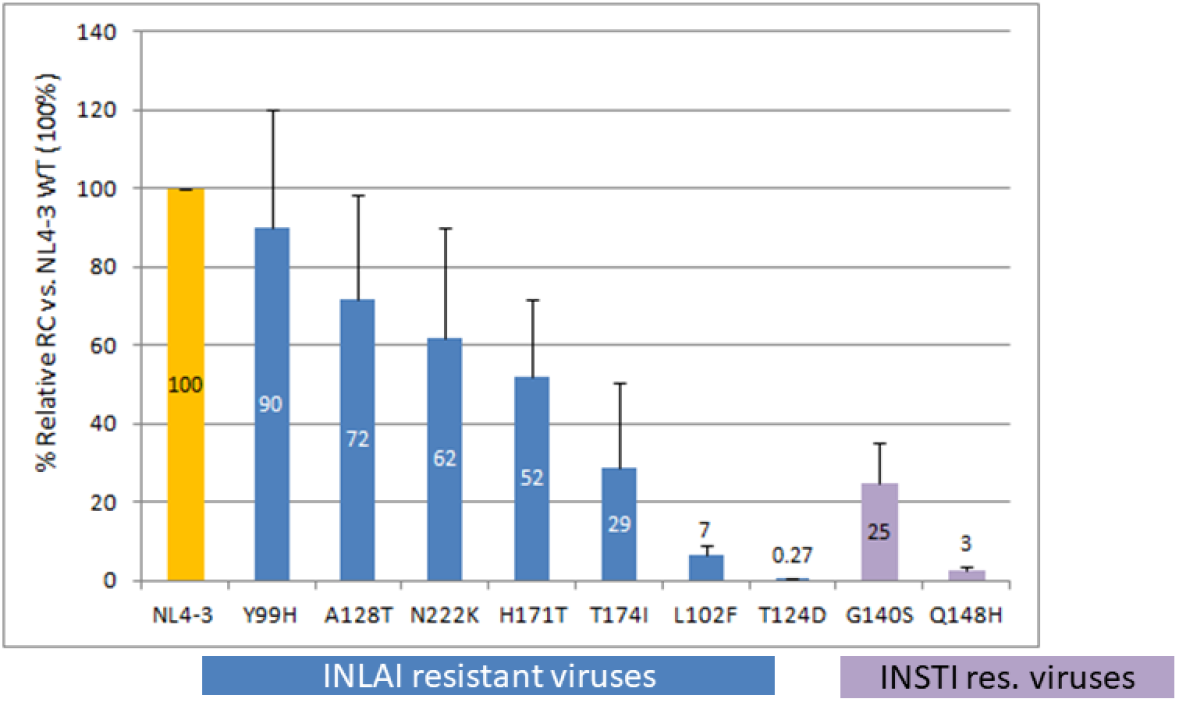

**S2 Table:** Statistics of data scaling and structure refinement.

| **Data collection: merging statistics** |  |
| --- | --- |
| Wavelength | 1.542 |
| Resolution range | 19.97 - 2.0 (2.072 - 2.0) |
| Space group | P 31 2 1 |
| Unit cell | 72.57 72.57 65.965 90 90 120 |
| Total reflections | 131582 (13489) |
| Unique reflections | 13873 (1364) |
| Multiplicity | 9.5 (9.9) |
| Completeness (%) | 99.19 (100.00) |
| Mean I/sigma(I) | 57.43 (11.20) |
| Wilson B-factor | 29.82 |
| R-merge | 0.02859 (0.1985) |
| R-meas | 0.03034 (0.2094) |
| R-pim | 0.009974 (0.06627) |
| CC1/2 | 1 (0.986) |
| CC* | 1 (0.997) |
| **Refinement statistics** |  |
| Reflections used in refinement | 13873 (1364) |
| Reflections used for R-free | 1386 (137) |
| R-work | 0.1955 (0.2156) |
| R-free | 0.2262 (0.2755) |
| CC(work) | 0.958 (0.896) |
| CC(free) | 0.942 (0.861) |
| Number of non-hydrogen atoms | 1157 |
| macromolecules | 1020 |
| ligands | 35 |
| solvent | 102 |
| Protein residues | 131 |
| RMS(bonds) | 0.008 |
| RMS(angles) | 1.03 |
| Ramachandran favored (%) | 98.32 |
| Ramachandran allowed (%) | 1.68 |
| Ramachandran outliers (%) | 0.00 |
| Rotamer outliers (%) | 0.00 |
| Clashscore | 3.85 |
| Average B-factor | 31.65 |
| macromolecules | 30.87 |
| ligands | 30.06 |
| solvent | 39.97 |

